# MYC Overexpression Drives Immune Evasion in Human Cancer that is Reversible Through Restoration of Pro-Inflammatory Macrophages

**DOI:** 10.1101/2022.05.13.491873

**Authors:** Renumathy Dhanasekaran, Aida S. Hansen, Jangho Park, Ian Lai, Nia Adeniji, Sibu Kuruvilla, Akanksha Suresh, Varsha Swamy, Dean W. Felsher

## Abstract

Cancers evade immune surveillance that in some, but not in many, cases can be reversed through immune checkpoint therapy. Here we report that the MYC oncogene suppresses immune surveillance, activates immune checkpoint expression, and predicts responsiveness to immune checkpoint inhibition. First, when MYC is genomically amplified and overexpressed in 33 different human cancers, this increases immune checkpoint expression, drives immune checkpoint therapeutic resistance, and is associated with both Th2-like immune profile, and reduced CD8 T cell infiltration. Second, experimentally, MYC-driven tumors suppress pro-inflammatory antigen-presenting macrophages with increased CD40 and MHCII expression, which in turn impedes T cell response. This MYC-driven suppression of macrophages can be reversed by combined but not individual blockade of PDL1 and CTLA4. Third, the depletion of macrophages abrogated the anti-neoplastic effects of PDL1 and CTLA4 blockade. Hence, MYC is a predictor of immune checkpoint responsiveness and a key driver of immune evasion through the suppression of pro-inflammatory macrophages. The immune evasion by MYC can be overcome by combined PDL1 and CTLA4 blockade.

**Statement of Significance:** MYC is the most commonly activated oncogene in human cancers. In this study, we identify macrophage-mediated immune evasion as a major therapeutic vulnerability of MYC-driven cancers. Our results have implications for developing effective immunotherapies for MYC-driven human cancers and also for prioritizing patients with MYC-driven tumors for combination immunotherapy.

## INTRODUCTION

The MYC oncogene is one of the most common drivers of human neoplasia, with more than 70% of human cancers having genetic alterations either in MYC or in one of the members of the proximal MYC network(1). This makes MYC a very desirable therapeutic target. However, no existing therapies directly target MYC. MYC contributes to tumorigenesis both by causing tumor intrinsic effects such as autonomous cellular proliferation, as well as by tumor extrinsic effects such as remodeling the tumor microenvironment via immune evasion (2). MYC inhibits anti-tumor immunity by multiple mechanisms, including impeding CD4+ T cell activation, promoting expression of immune checkpoints like PDL1 and CD47, and blocking natural killer (NK) cell activation (2–7). These immune effects suggest that MYC-driven tumors may be vulnerable to specific combinations of immune therapies.

MYC activation is a major oncogenic driver of many types of human cancer, including cancers of epithelial origin such as hepatocellular carcinoma (HCC), breast cell cancer, and colorectal carcinoma. (8–12) HCC often has a dismal prognosis. (13,14) Immune checkpoint inhibitors (ICI) have shown promise in many human cancers but, notably, have failed as first-line monotherapy for HCC. (15,16). On the other hand, the combination of PD1 AND CTLA4 blockade has shown promise in a subset of patients (NCT03298451)(17). Understanding the mechanisms by which individual or combined ICIs are effective in subsets of patients could enhance their efficacy.

We hypothesize that MYC is a major driver of immune evasion and dictates immune checkpoint therapeutic response. Here we demonstrate the combination of immune checkpoint inhibitors PDL1 and CDL4 can overcome MYC immune evasion by restoring macrophage activation via CD40 and MHCII expression, thereby eliciting robust anti-tumor CD8+ T-cell responses.

## RESULTS

### The MYC Oncogene Drives Human Tumorigenesis Associated with Immune Evasion

We evaluated the role of *MYC* in the immune status of human cancers using the cancer genome atlas (TCGA) pan-cancer studies (n= 11,069 samples, n=33 cancers, Fig 1A). First, *MYC* was most commonly activated by genomic amplification (n=889, 8%), especially in solid tumors like HCC (11%) (Fig 1A; Supp Fig 1A)(18). *MYC* amplification was strongly associated with *MYC* gene overexpression (p<0.0001; Supp Fig 1B). Even tumors without *MYC* amplification showed overexpression of the gene signature associated with *MYC* amplification (Fig 1A), highlighting the ubiquitous nature of MYC activation in cancers. Both *MYC* amplification and *MYC* mRNA overexpression were associated with worse overall and progression-free survival (Fig 1B). Further, *MYC*-amplified tumors had higher tumor grade (p= 7.0 × 10^−8^) and belonged to more advanced stages (p=2.7 × 10^−7^) (Supp 1C-D). Thus, MYC activation by amplification and/or overexpression is a global driver of tumorigenesis in human cancers.

**Figure 1.**
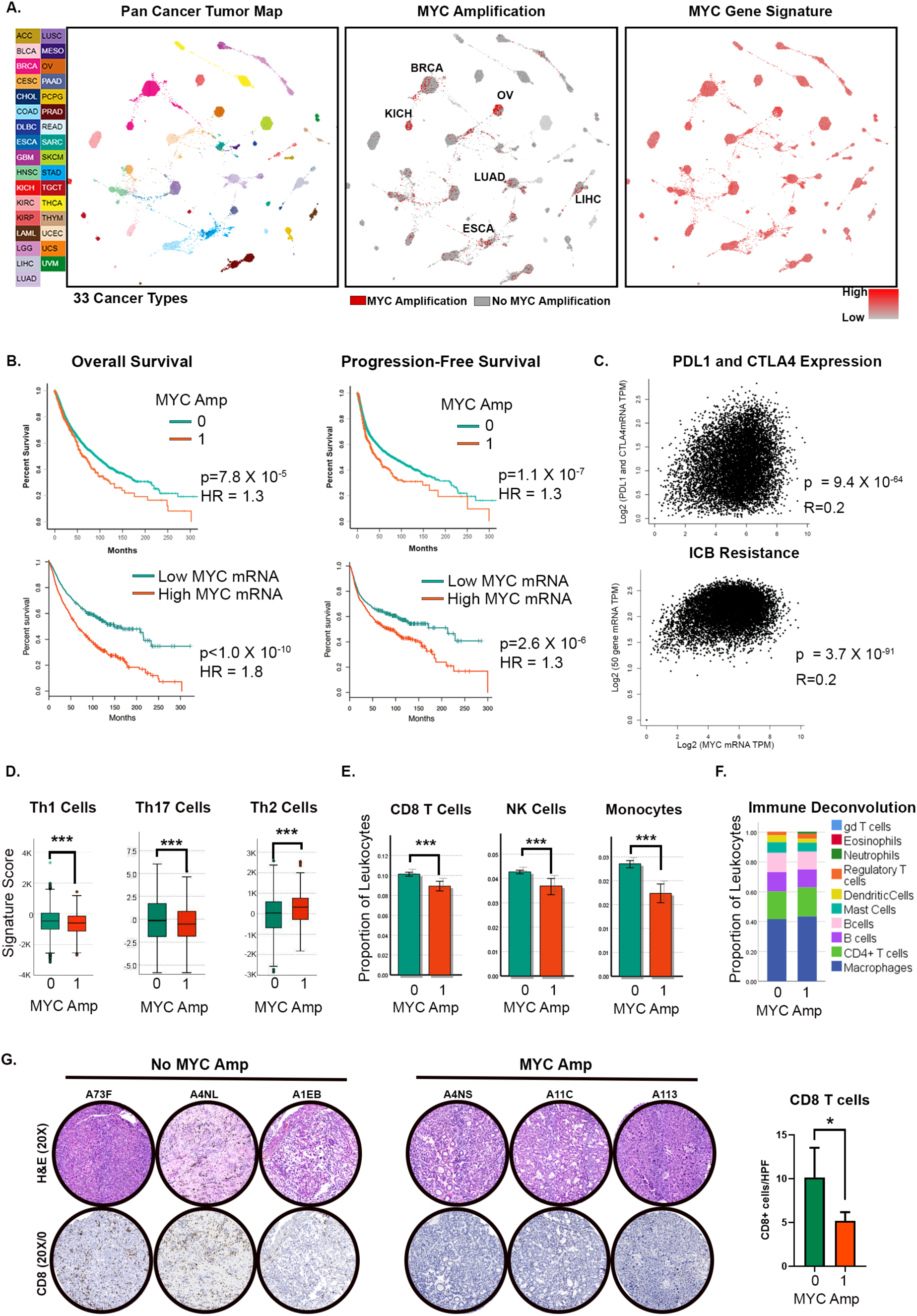
The MYC Oncogene Drives Human Tumorigenesis Associated with Immune Evasion. A. TumorMap rendering of the Pan-Cancer-Atlas of 33 cancers; tumors are defined by different tissues of origin in the first box. Second box shows the prevalence of *MYC* genomic amplification across cancers. Epithelial tumors with high rates of *MYC* amplification are annotated. Third box shows the gene expression of a signature derived from *MYC* amplification. (LAML- Acute Myeloid Leukemia, ACC- Adrenocortical carcinoma, BLCA- Bladder Urothelial Carcinoma, LGG- Brain Lower Grade Glioma, BRCA- Breast invasive carcinoma, CESC- Cervical squamous cell carcinoma and endocervical adenocarcinoma, CHOL- Cholangiocarcinoma, LCML- Chronic Myelogenous Leukemia, COAD- Colon adenocarcinoma, ESCA- Esophageal carcinoma, GBM- Glioblastoma multiforme, HNSC- Head and Neck squamous cell carcinoma, KICH- Kidney Chromophobe, KIRC- Kidney renal clear cell carcinoma, KIRP- Kidney renal papillary cell carcinoma, LIHC- Liver hepatocellular carcinoma, LUAD- Lung adenocarcinoma, LUSC- Lung squamous cell carcinoma, DLBC- Lymphoid Neoplasm Diffuse Large B-cell Lymphoma, MESO- Mesothelioma, OV- Ovarian serous cystadenocarcinoma, PAAD- Pancreatic adenocarcinoma, PCPG- Pheochromocytoma and Paraganglioma, PRAD- Prostate adenocarcinoma, READ- Rectum adenocarcinoma, SARC- Sarcoma, SKCM- Skin Cutaneous Melanoma, STAD- Stomach adenocarcinoma, TGCT- Testicular Germ Cell Tumors, THYM- Thymoma, THCA- Thyroid carcinoma, UCS- Uterine Carcinosarcoma, UCEC- Uterine Corpus Endometrial Carcinoma, UVM- Uveal Melanoma). B. Overall and progression-free survival in patients with *MYC* amplification and *MYC* overexpression (top and bottom quartiles). C. Correlation of *MYC* expression with gene expression of immune checkpoints *PDL1* and *CTLA4* (top) and with gene signature of immune checkpoint blockade (ICB) (Jiang et al. 2018; bottom). D. Expression of gene signatures associated with T helper cells 1 (Th1), Th17 and Th2 compared between tumors with and without MYC amplification. E. Expression of gene signatures associated with CD8 T cells, NK cells and monocytes compared between tumors with and without MYC amplification. F. Immune deconvolution analysis showing comparison of abundance of different immune subsets between tumors with and without MYC amplification. G. Immunohistochemistry showing CD8 T-cell infiltration in representative HCC tumors from the TCGA cohort with (n=20) and without MYC amplification (n=8). *p<0.05, ***p<0.0001

Next, we found that MYC activation influences the immune state of human cancers. First, the expression of MYC strongly correlated with the expression of the immune checkpoints *PDL1* and *CTLA4* (p=9.4×10^−64^) (Fig 1C). Second, MYC overexpression was associated with a gene signature predictive of resistance to immune checkpoint blockade (19) (9=3.7×10^−91^) (Fig 1C). Third, MYC-amplified tumors had lower expression of anti-tumor Th1 and Th17 signatures and higher expression of pro-tumorigenic Th2 immune signature (p<0.001) (Fig 1D). Fourth, MYC-amplified tumors had lower infiltration with anti-tumor immune subsets like CD8+ T cells, NK cells, and monocytes (all p<0.001), with no major differences in other subsets upon immune deconvolution analysis (Fig 1E-F). Finally, we examined if MYC overexpression in HCC was associated with immune suppressiveness since MYC is frequently amplified and overexpressed in HCC (Supp Fig 2A-B) and HCC exhibits limited responsiveness to immune checkpoint blockade.(15,20) Indeed, by IHC, we found that MYC amplification was associated with lower CD8+ T cell infiltration (Fig 1G). Thus, MYC overexpression is common in human cancer and is associated with a poor prognosis. Moreover, MYC-driven tumors are immune evasive and less likely to respond to immune checkpoint blockade.

### MYC causes Reversible Suppression of Anti-Tumor Immune Response

To examine how MYC suppresses the immune response, we used a MYC-driven conditional transgenic mouse model of HCC (21). The transcriptional profile of our transgenic mouse model of MYC-HCC was similar to the transcriptome to multiple human HCC cohorts including ten human HCC datasets and the TCGA data (similarity z score 27.2, p value= 1×10^−15^) (Fig 2A). Specifically, there was an overlap between mouse MYC-HCC transcriptome and the human HCC immune signatures of inactivated 41BB-ve PD1+ T cells in human HCC (23) (similarity z-score 3.2, p value= 1×10^−9^) and exhausted PD1+ CD8 T cells (22) (similarity z-score 16.7, p= 1×10^−21^) (Fig 2B). Canonical pathways and biological functions of anti-tumor immunity were significantly downregulated in both our murine MYC-HCC model and human HCC (Fig 2B).

**Figure 2:**
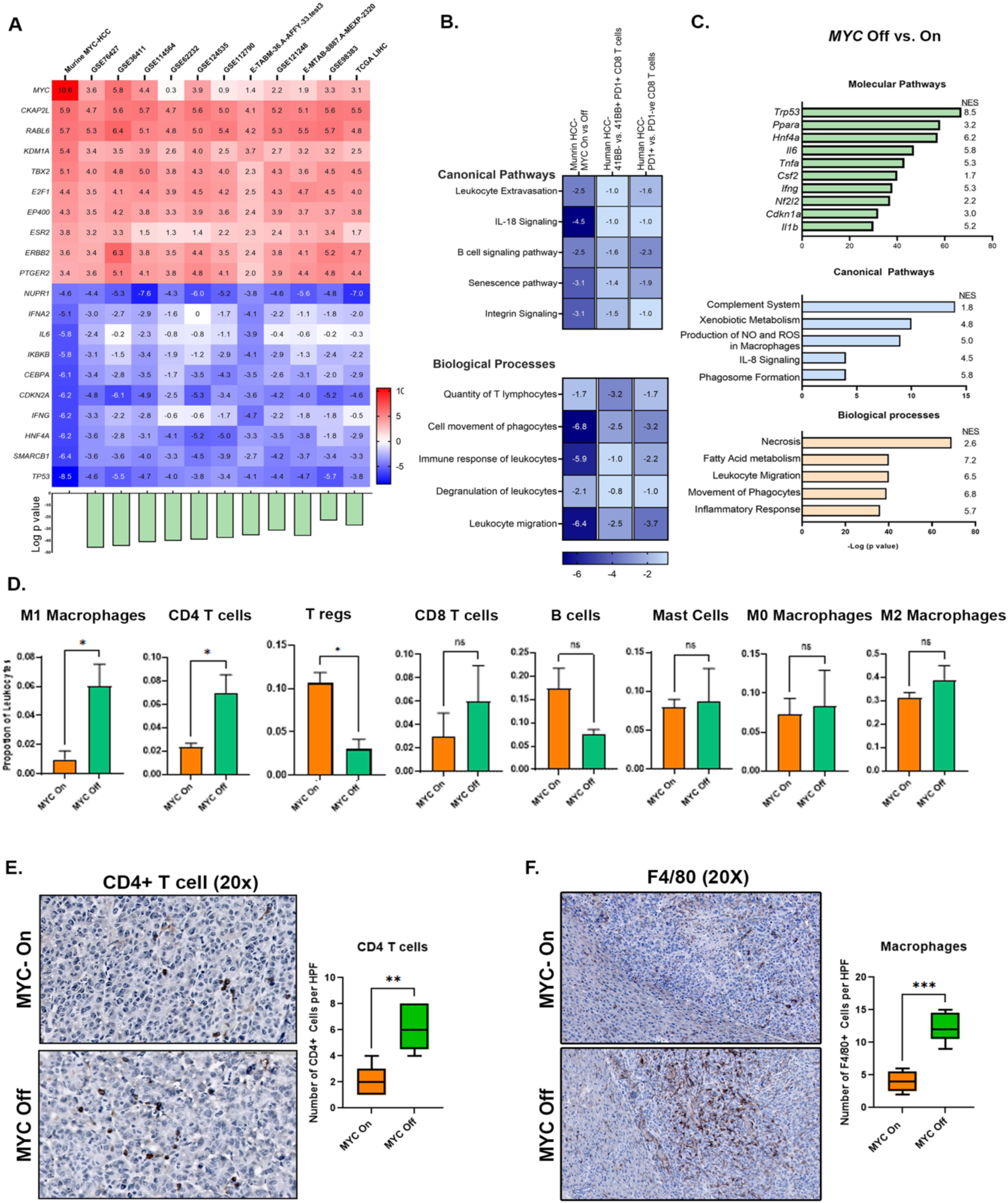
MYC causes Reversible Suppression of Anti-Tumor Immune Responses. A. Heat map shows similarity between MYC-driven murine HCC and 11 human HCC transcriptome data sets including the TCGA cohort; the top 10 upregulated and downregulated genes with enrichment z-scores across the different data sets are shown. Barplot at the bottom shows log of *p-*value of similarity between the individual cohorts and murine MYC-HCC. B. Heatmap shows z-scores of enrichment of canonical pathways and biological functions in murine MYC-HCC transcriptome and the human HCC immune signatures of exhausted PD1+ CD8 T cells ^22^ and inactivated 41BB-ve PD1+ T cells in human HCC ^23^. C. Top molecular pathways, canonical pathways and biological processes activated in murine MYC HCC upon MYC inactivation. D. Immune deconvolution analysis showing comparison of abundance of different immune subsets in murine MYC HCC upon MYC inactivation. E. Immunohistochemistry shows the abundance of CD4 T-cells upon MYC inactivation. E. Immunohistochemistry shows the abundance of macrophages upon MYC inactivation. **p<0.01, ***p<0.0001

Next, we examined the functional changes induced upon MYC inactivation in the mouse model of MYC-driven HCC (MYC On vs. Off DE genes 728 upregulated, 1352 downregulated; pAdj<0.05). Inactivating MYC in the MYC-HCC resulted in the upregulation of anti-tumor immune pathways including the *Il6, Tnfa* and *Ifng* pathways, and activation of inflammatory biological processes like phagosome formation, production of ROS by macrophages, and phagocyte migration (Fig 2C). Additionally, MYC inactivation led to the recruitment of pro-inflammatory M1-macrophages and CD4+ T cells, and a decrease in the proportion of pro-tumorigenic immune cells like regulatory T (Treg) cells (Fig 2D). These changes were confirmed by IHC (Fig 2E-F). Thus, the mouse MYC-HCC and human HCC exhibit similar immune profiles associated with exhausted T cells. Further, in the mouse model, MYC inactivation reverses immune evasion by activating innate and adaptive anti-tumor immunity.

### Combined Anti-PDL1 and Anti-CTLA4 overcome MYC induced immune evasion

We evaluated if inhibiting the immune checkpoints PDL1 and/or CTLA4 could overcome MYC-induced immune evasion (Fig 3A). In mouse MYC-HCC, only mice treated with both anti-PDL1 and anti-CTLA4, but not either alone, showed significantly delayed progression on three-dimensional tumor volume assessment by MRI compared to mice treated with control antibody (mean tumor volume [858.31 μm^3^, SEM = 53.60] vs. [1897.67 μm^3^, SEM = 258.62]; p = 0.004)) (Fig 3B-C). Mice treated with combined PDL1 and CTLA4 blockade had smaller and fewer liver tumors than mice treated with control antibody ([858.31 μm^3^, SEM = 53.60] vs. [1897.67 μm^3^, SEM = 258.62]; p = 0.004) (Fig 3D-E). However, there were no differences in final tumor volumes of mice treated with anti-PDL1 (1235.81 μm^3^, SEM = 288.87) or anti-CTLA4 alone (1922.64 μm^3^, SEM = 509.45) compared to IgG treated mice (p = 0.126; p = 0.966 respectively) (Fig 3D-E). The combination therapy led to a decrease in the proliferative index of the tumor as measured by phospho-histone 3 staining without affecting MYC expression levels in the tumor cells (Supp Fig 3). Thus, only combined, but not individual, inhibition of PDL1 and CTLA4, impeded tumor progression.

**Figure 3.**
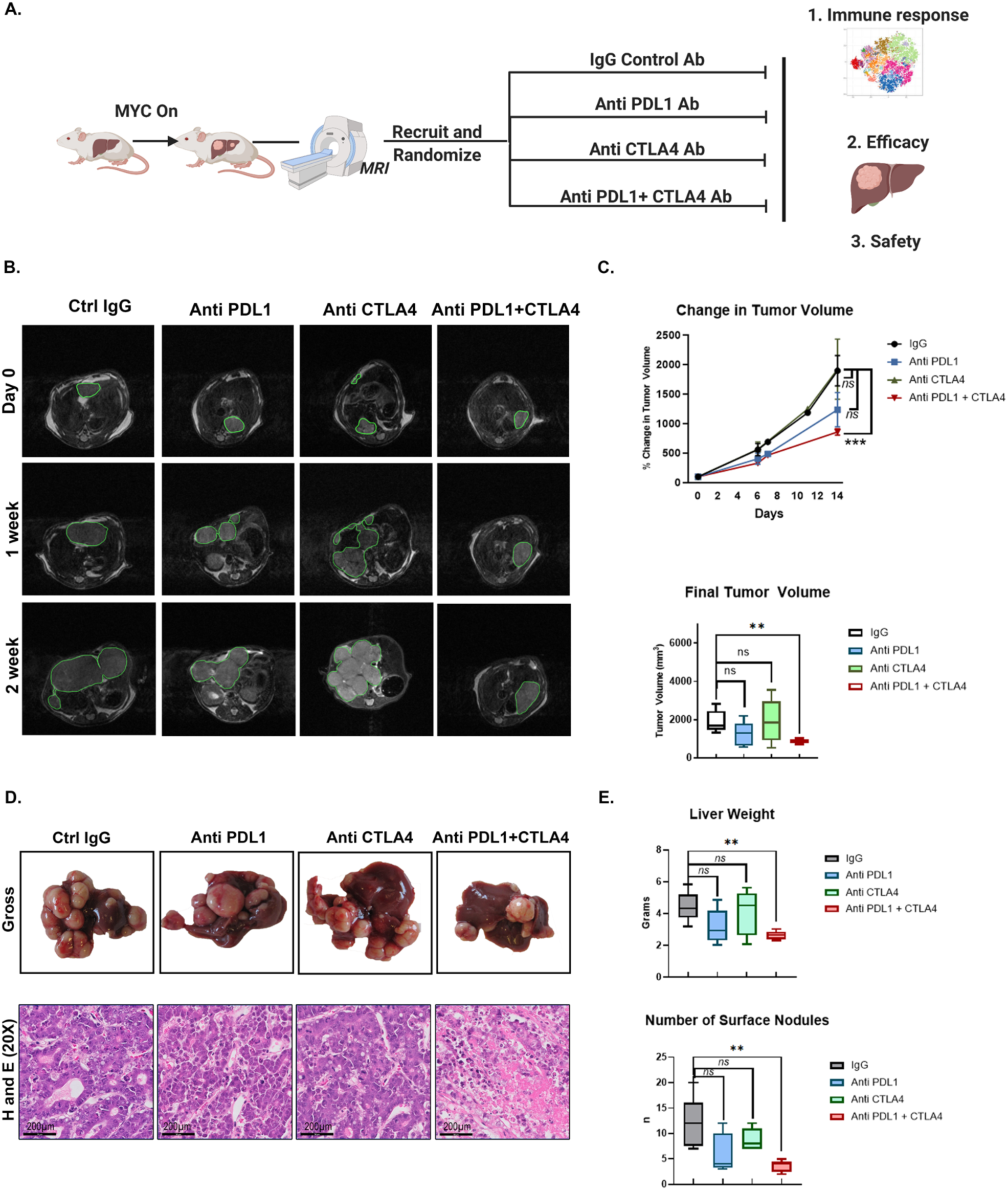
Combined Anti- PDL1 and Anti-CTLA4 delays tumor progression in MYC-HCC. A. Experimental scheme of MYC-HCC treatment with IgG control, PDL1 antibody, CTLA4 antibody, or their combination. B. Weekly MRI showing tumor progression in representative MYC-HCC mice in the 4 treatment groups. C. Quantification of volumetric tumor measurement using MRI of MYC-HCC mice in the 4 treatment groups. D. End-of-treatment gross appearance, histology of representative MYC-HCC mice in the 4 treatment groups. E. Quantification of liver tumor burden at end-of-treatment of MYC-HCC mice in the 4 treatment groups. **p<0.01, ***p<0.0001

Treatment with combination therapy did not result in weight loss or hepatotoxicity in mice (Fig 4A). End-of-treatment analysis revealed slight lymphopenia with individual or combined treatments. Treatment with anti-CTLA4 alone and in combination with anti-PDL1 resulted in lower peripheral blood monocyte count (Fig 4B). Histological evaluation of liver, colon, and lungs by a blinded pathologist showed no areas of severe autoimmune organ damage with the combination therapy even though immunohistochemical analysis revealed higher levels of CD8+ T cell infiltration with combination therapy (Fig 4C). Thus, the combination of anti-PDL1 and anti-CTLA4 shows synergistic efficacy in overcoming MYC-induced immune evasion without causing severe toxicity.

**Figure 4.**
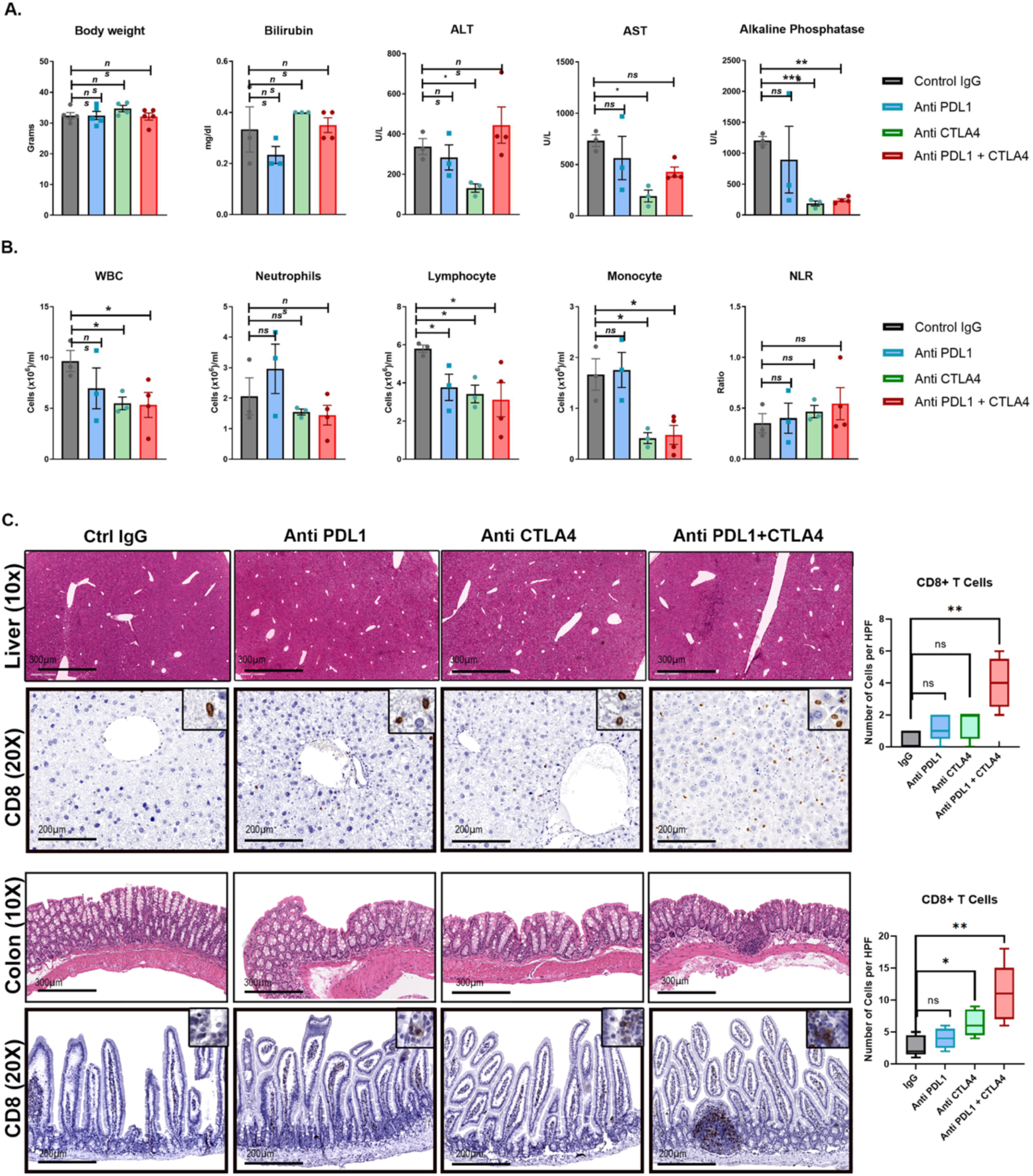
Safety profile of combined Anti- PDL1 and Anti-CTLA in MYC-HCC. A. Body weight and liver tests at end-of-treatment of MYC-HCC bearing mice treated with either IgG control or PDL1 antibody or CTLA4 antibody or their combination. B. Peripheral white cell counts at end-of-treatment of MYC-HCC bearing mice treated with either IgG control or PDL1 antibody or CTLA4 antibody or their combination. C. Histology and CD8 T cell infiltration in normal surrounding liver and colon tissue of mice treated with the 4 treatment groups. *p<0.05, **p<0.01

### Dual targeting of PDL1 and CTLA4 Restores Macrophage-Mediated Anti-tumor immunity

Next, we evaluated how dual targeting of anti-PDL1 and CTLA4 remodeled the immune microenvironment of the tumors. IHC analysis showed that the combination of PDL1 and CTLA4 blockade elicited recruitment of a higher number of CD4+ T cells (p=0.005) and CD8+ T cells (p<0.0001) than individual monotherapies (Fig 5A). Interestingly, we observed significantly increased macrophage infiltration in tumors treated with the combination of PDL1 and CTLA4 compared to IgG treated mice (p=0.005) but not in mice treated with anti- PDL1 (p=0.31) or anti- CTLA4 (p=0.06) as monotherapy (Fig 5B). Despite an overall increase in macrophage infiltration, the proportion of immunosuppressive PDL1+ macrophages was lower in mice treated with combined anti-PDL1 and anti CTLA4 compared to mice treated with IgG control antibody (p=0.001) (Supp Fig 4). Thus, dual targeting of PDL1 and CTLA4 elicits a more robust T cell immune response and macrophage infiltration than either monotherapy.

**Figure 5.**
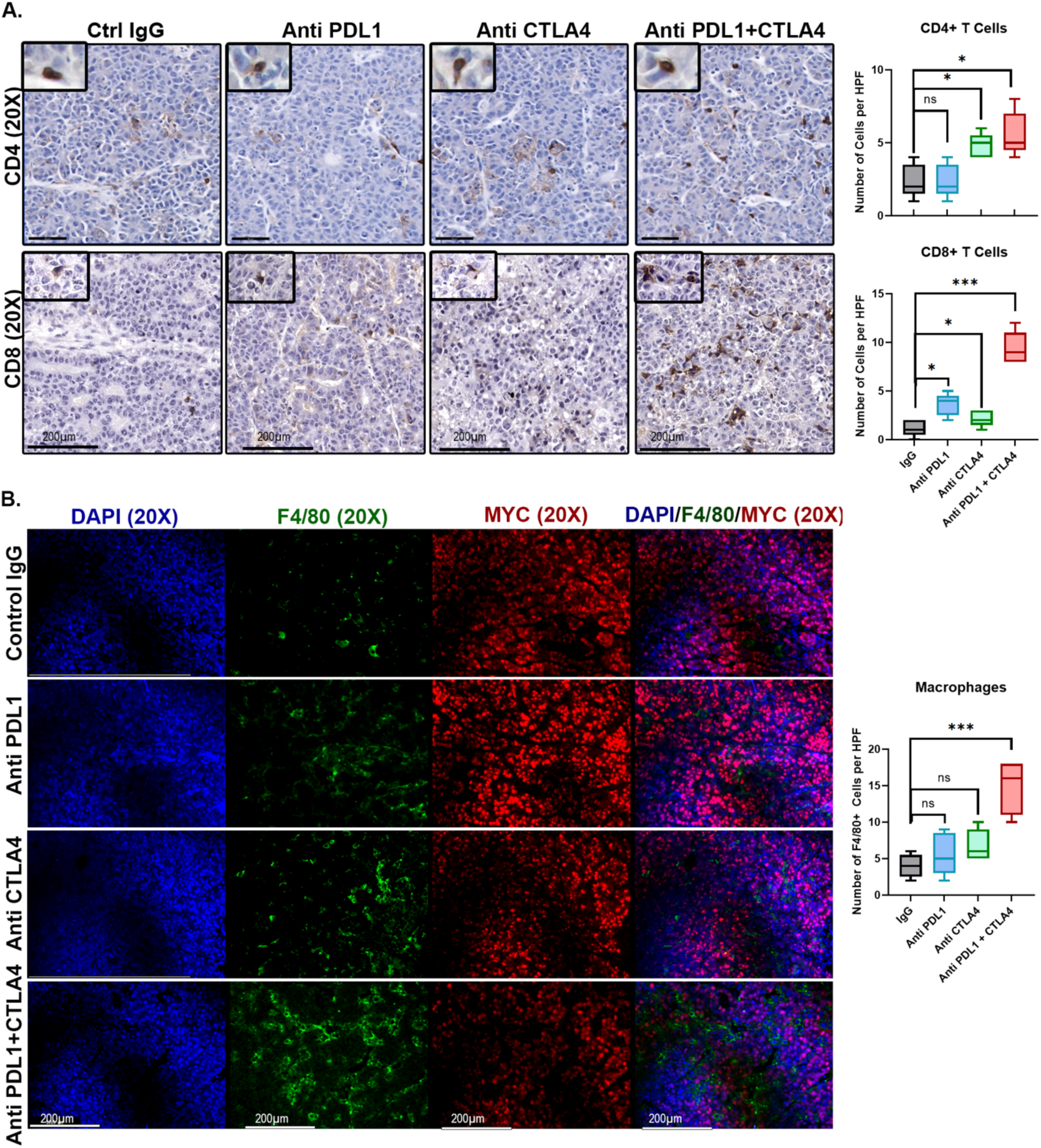
Dual targeting of PDL1 and CTLA4 Restores Macrophage-Mediated Anti-tumor immunity. A. Immunohistochemistry for CD4 T cells and CD8 T cells in MYC-HCC bearing mice treated with IgG control, PDL1 antibody, CTLA4 antibody, or their combination. Boxplots show quantification of cell counts. B. Immunofluorescence for F4/80 and MYC in MYC-HCC-bearing mice treated with IgG control, PDL1 antibody, CTLA4 antibody, or their combination. Boxplots show quantification of cell counts. *p<0.05, ***p<0.0001.

### Combined PD-L1 and CTLA-4 Inhibition Elicits Early Repolarization of Myeloid Cells

To determine the sequence of macrophage and T cell activation with immune checkpoint inhibition, we evaluated immune responses at an earlier time point of 1 week after treatment with PDL1 and/or CTLA4 inhibitors. We used high-dimensional single-cell profiling of the tumor immune infiltrates with mass cytometry (CyTOF) (Fig 6A). After dimensionality reduction of the high-order parametric data, twenty-seven unique immune subsets were identified (Fig 6B and Supp Fig 5A). The most significant changes in the mice treated with combined anti-PDL1 and anti-CTLA4, compared to the individual monotherapies, were found in the myeloid compartment (Fig 6C). The frequency of pro-inflammatory macrophages (Ly6C^High^/CCR2^High^) was significantly increased, and immunosuppressive macrophages (PDL1+/Ly6C^Low^/CCR^Low^) was significantly decreased (Fig 6D) in the tumors treated with both anti-PDL1 and anti-CTLA4, and not either alone, compared to mice treated with IgG control (Fig 6D).

**Figure 6.**
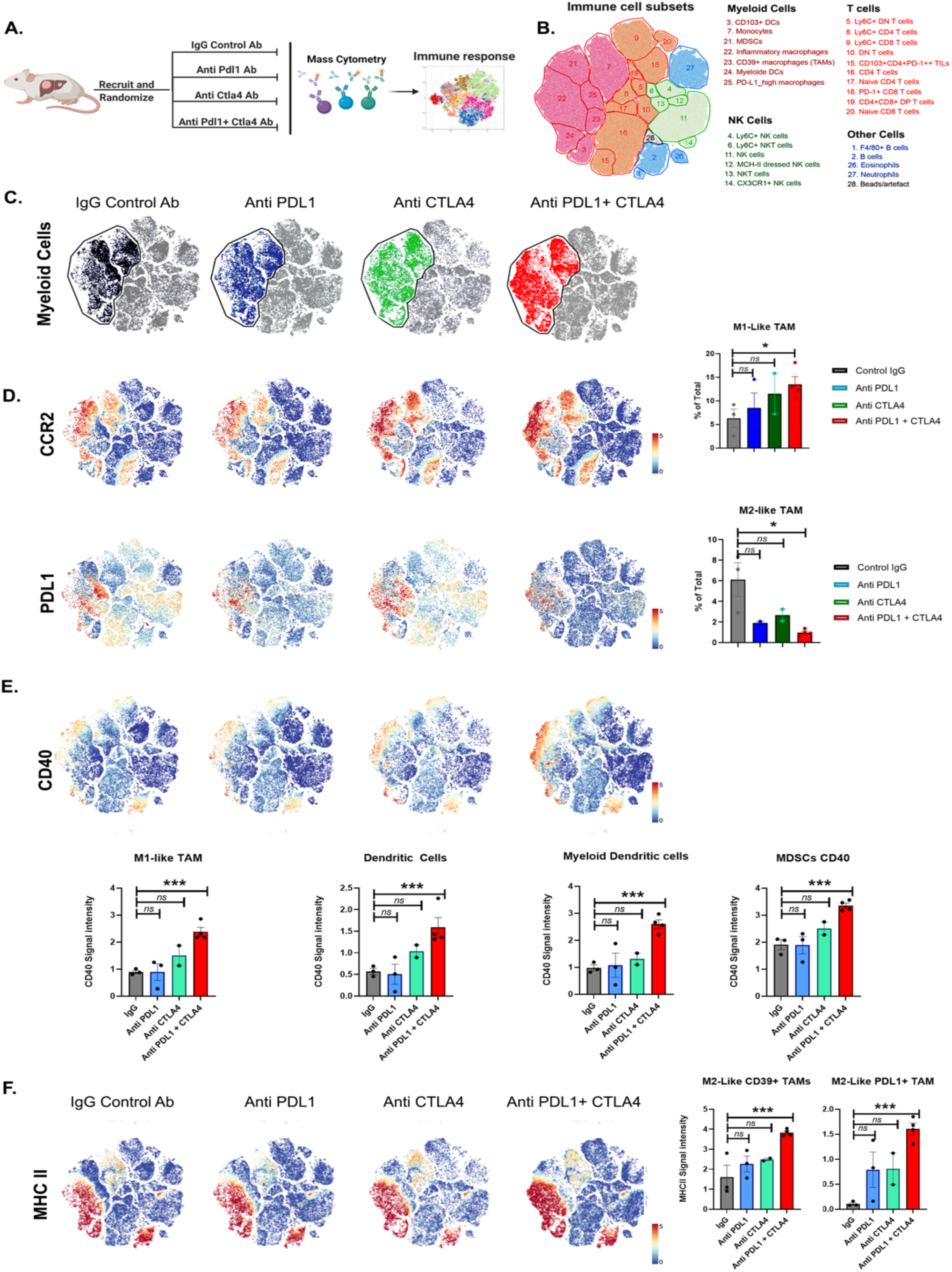
Mass cytometry reveals unique early immune response to dual targeting of PDL1 and CTLA4. A. Experimental scheme of MYC-HCC treatment with IgG control, PDL1 antibody, CTLA4 antibody, or their combination for 1 week, followed by mass cytometry analysis. B. tSNE plot grouped by major immune subsets identified in the tumor immune microenvironment. C. tSNE plots showing CCR2 and PDL1 expression in the MYC-HCC bearing mice treated with IgG control, PDL1 antibody, CTLA4 antibody, or their combination. Boxplot shows quantification of CCR2+ Ly6^High^ M1-Like Macrophages and PDL1+ M2-Like Macrophages respectively. D. tSNE plots showing CD40 expression in the MYC-HCC bearing mice treated with IgG control, PDL1 antibody, CTLA4 antibody, or their combination. Boxplot shows quantification of CD40 expression in different myeloid subsets. E. tSNE plots showing MHCII expression in the MYC-HCC bearing mice treated with IgG control, PDL1 antibody, CTLA4 antibody, or their combination. Boxplot shows quantification of MHCII in CD39+ and PDL1+ M2-Like Macrophages.

CD40 expression on myeloid cells is known to activate the tumoricidal activities of macrophages (24,25). The combination therapy, but not either anti PDL1 or anti CTLA4 alone, significantly increased CD40 expression on multiple myeloid cells including inflammatory macrophages (Ly6C^High^/CCR2^High^), conventional dendritic cells (CD11c+/CD11b-/CD103+), myeloid dendritic cells (CD11c+/CD11b+/CD103-) and MDSCs (Fig 6E). On the other hand, immunosuppressive macrophage subsets like the PDL1+ TAMs and CD39+ TAMs showed significantly increased expression of MHC II with combination anti-PDL1 and anti-CTLA4, but not with the individual monotherapies (Fig 6F). We did find a trend towards increased effector CD8+ T cells (LyC^High^/PD1^Low^) with combination therapy but no other significant differences in the proportion of T cell subpopulations at this early stage of treatment (Supp Fig 5C). There were no significant changes in other innate immune cells like NK cells, eosinophils, or neutrophils (Supp Fig 5B).

Thus, we show that combined blockade of both PDL1 and CTLA4, and not either alone, elicits an early immune response with the expansion of the pro-inflammatory myeloid compartments and repolarization to an anti-tumor profile with upregulation of CD40 and MHC-II.

### Macrophages are Essential for the anti-tumor efficacy of Immune Checkpoint Therapy

To evaluate the causal role of macrophages, we depleted macrophages prior to treatment with combined PDL1 and CTLA4 blockade (Fig 7A). We used the anti-CSF1R antibodies to deplete macrophages (26,27), after first confirming that the CSF1R antibody effectively depleted macrophages, without impacting tumor progression in MYC-driven HCC (Supp Fig 6A-C).

**Figure 7.**
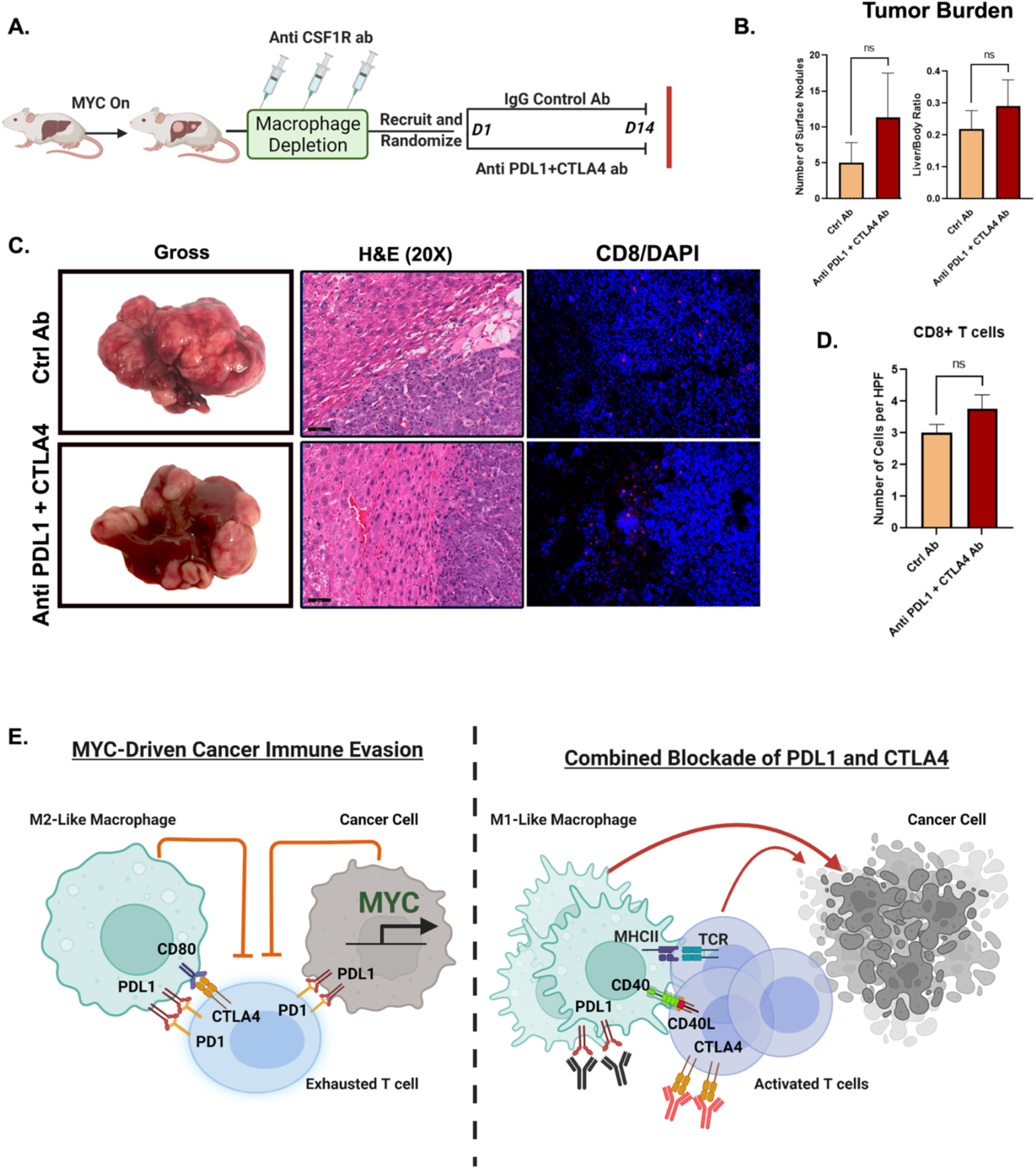
Macrophages are Essential for the anti-tumor efficacy of Combined Immune Checkpoint Therapy. A. Experimental scheme of macrophage depletion in MYC-HCC followed by treatment with either IgG control or dual PDL1 and CTLA4 antibodies. B. Quantification of tumor burden at end-of-treatment with either IgG control or dual PDL1 and CTLA4 antibodies in macrophage-depleted MYC-HCC mice. C. End-of-treatment gross appearance, histology and IHC for F4/80 confirming macrophage depletion in representative MYC-HCC treated with IgG control or dual PDL1 and CTLA4 antibodies. D. Immunofluorescence for CD8 T cells in representative macrophage-depleted MYC-HCC-bearing mice treated with IgG control or dual PDL1 and CTLA4 antibodies. E. Figurative representation of the mechanism of efficacy of combined PDL1 and CTLA4 therapy in MYC-HCC. MYC-driven cancers are immune evasive with PDL1 overexpression on cancer cells and macrophages which in turn lead to T cell exhaustion. Treatment with CTLA4 and PDL1 inhibitors leads to repolarization of macrophages to the M1-like phenotype with increased expression of CD40 and MHCII, which leads to enhanced antigen presentation and robust T cell activation resulting in delayed tumor progression.

There was no difference in liver tumor burden between macrophage-depleted mice treated with combination PDL1 and CTLA4 inhibition or control antibody (liver/body ratio 0.25 vs. 0.17, p=0.23) (Fig 7B). Further, we confirmed that the combined blockade of PDL1 and CTLA4 was not able to elicit antitumor CD8 T cell response in macrophage-depleted mice (Fig 7C-D). Together, this shows that macrophage depletion blocks the anti-tumor efficacy of combined PDL1 and CTLA4 inhibition. Thus, our data demonstrate that combined PDL1 and CTLA4 blockade requires the activation of macrophages to overcome MYC-induced immune evasion (Fig 7E).

## DISCUSSION

We show that MYC is causally involved in immune evasion during tumorigenesis which can be overcome through combination immune checkpoint blockade. A pan-cancer analysis of 33 human tumors reveals that MYC is not only commonly amplified and overexpressed, but it is also associated with increased immune checkpoint expression, immunotherapy resistance, a Th2-like immune profile, and reduced anti-tumor CD8 T-cell infiltration. We demonstrate that MYC is causally involved in blocking immune checkpoint responses in a murine model of MYC-driven HCC. Notably, MYC immune evasion can be abrogated through combined immune checkpoint blockade with anti-PDL1 and CTLA4, which was associated with the causally required activation of macrophages to a pro-inflammatory phenotype. The restoration of immune responsiveness was associated with increased expression of CD40 and MHCII on antigen-presenting cells (APCs) and robust anti-tumor T cell responses (Fig 7E). We conclude that MYC is a key driver of immune evasiveness in human cancer, blocking both innate and adaptive antitumor immune responses. Further, we identify MYC-driven tumors as a specific subset of cancers that are unlikely to respond to monotherapy with immune checkpoint inhibitors and need combination immunotherapy to achieve tumor remission.

Inhibition of immune checkpoints like PDL1 or CTLA4 has shown promising efficacy in many cancers (28,29). However, less than a third of patients respond to ICI monotherapy, making it critical to understand mechanisms of resistance to immunotherapy. While the role of MYC in driving immune evasion has been demonstrated by previous studies, (4,5,30–32), its functional relevance in modulating response to immune checkpoint inhibition is not understood. Our study identifies MYC oncogene amplification or overexpression as a major driver of tumor-intrinsic resistance to immune checkpoint inhibition. We demonstrate that MYC-driven cancers exhibit primary resistance to monotherapy with PDL1 or CTLA4 blockade, implying that additional aspects of the immune response are modulated by MYC. These results imply that MYC-amplification or MYC-overexpression can potentially serve as a biomarker to prioritize patients to be considered for combination immunotherapies, rather than individual immune checkpoint inhibitors.

Our results identify a unique mechanistic basis for the synergistic efficacy of combined PDL1 and CTLA4 blockade, beyond just additive effects of their individual mechanisms. We suggest a model whereby the combination of anti-PDL1 and CTLA4 restores immune responsiveness by the activation of pro-inflammatory CCR2+ macrophages, which we show are causally required, as macrophage depletion abrogated the anti-tumor efficacy of PDL1 and CTLA4 blockade (Figure 7E). These macrophages appear to be involved in the subsequent recruitment of adaptive immune cells including CD4+ T cells and cytotoxic CD8+ T cells. We suggest that the upregulation of at least two ligands on APCs, CD40, and MHCII, is involved in the mechanism of activating adaptive immunity. CD40 expression is known to license APCs to activate cytotoxic T cells and invigorate the tumoricidal activity of macrophages (33). Increased MHCII expression enables efficient antigen presentation by APCs to CD4+ T cells (34). The upregulation of both CD40 and MHCII on APCs, seen uniquely with combined PDL1 and CTLA4 blockade and not with the monotherapies, could be a key mechanism that contributes to the anti-tumor efficacy of the combination.

Finally, no existing therapies that directly target the MYC oncogene exist (35,36). Our results suggest that in human tumors with MYC activation, combined PDL1 and CTLA4 inhibition may be effective in restoring macrophage-mediated anti-tumor immune response.

## METHODS

### Human pan-cancer TCGA analysis

We used the TumorMap (https://tumormap.ucsc.edu/), a tool that enables grouping the cancer genome atlas (TCGA) samples in a visually accessible way. Tumor Map uses dimensionality reduction methods to condense high-dimensional omics data to a two-dimensional space. In the TumorMap, each node is a sample, and clusters of samples indicate groups with similar genomic alteration events or oncogenic signatures. We used the TCGA pan-cancer data to visualize samples with MYC amplification (37). We visualized the MYC gene signature derived from gene set enrichment analysis of MYC amplification across the TCGA pan-cancer samples (http://www.broadinstitute.org/gsea/msigdb/cards/chr8q24). The Spearman test was used for correlation analysis. Kaplan-Meier analysis was performed for survival analysis. We used the immune signatures and immune deconvolution analysis published in the pan-cancer TCGA immune landscape data (38).

### RNA sequencing

RNA sequencing of MYC-HCC was performed at the Beijing Genomics Institute (BGI) using their BGIseq 500 platform single end 150 bp, 20 million reads per sample. MYC-activated tumors (MYC-On, n=3)) and tumors four days after MYC inactivation (MYC-Off, n=3) underwent whole transcriptome sequencing. Gene expression level was quantified by a software package called RSEM. For each RNA sequencing data sample deposited in Gene Expression Omnibus (GEO), we counted the number of identified expressed genes and calculated its proportion to the total gene number in the database. DEseq software was used to perform differential expression analysis. Ingenuity Pathway Analysis (IPA, Qiagen) was used to perform functional pathway analysis and similarity analysis. The CIBERSORT gene expression deconvolution package was used to estimate the immune cell composition in the MYC-HCC (39). The LM22 signature was used as the immune cell gene signature after the careful modification of genes in the signature to their respective mouse orthologs. The settings for the run were: 1000 permutations with quantile normalization disabled. The Student’s t-test was used to infer the statistical significance of the predicted immune cell populations where p<0.05 was considered significant.

### Transgenic Mice

Animals were housed in a pathogen-free environment at Stanford University and all procedures were performed in accordance with Stanford’s Administrative Panel on Laboratory Animal Care (APLAC) protocols. LAP-tTA/tet-O-MYC transgenic lines were used, as previously described (21). Mice were administered weekly doses of 0.1 mg/mL doxycycline (Sigma) in drinking water during mating and until four weeks of age. At 4 weeks, mice were taken off doxycycline. Mice were screened for tumors via MRI at approximately 2-3 months of age, at which time they developed tumors between 50-150mm^3^.

### Magnetic Resonance Imaging (MRI)

MRI was performed using a 7T small animal MRI (Agilent conversion) with a 40 mm Varian Millipede RF coil (ExtendMR LLC, Milpitas, CA) at the Stanford Small Animal Imaging Facility as previously described (40,41). Briefly, animals were anesthetized with 1-3% isoflurane and placed into the MRI scanner containing a 40 mm Varian Millipede RF coil (ExtendMR LLC, Milpitas, CA). ParaVision (PV6.01) was used to acquire the DICOM images, and tumor volumes were quantified from images using Osirix image processing software (Osirix, UCLA, and Los Angeles, CA). Tumor sizes in mice were monitored weekly as well as once before and after the completion of treatments.

### Immunotherapy Treatment

Two days following tumor detection, mice were enrolled into one of four treatment groups. Control Rat IgG (BioXCell) and αPDL1 (clone 10 F.9G2, BioXCell) antibodies were given I.P. (100 μg/mouse) every other day, αCTLA4 antibody (clone UC10-4 F10-11, BioXCell) was given i.p. (100 μg/mouse) every three days twice. For the long-term treatment, mice were treated for two weeks, and for the mass cytometry experiment, mice were treated for one week. For the macrophage-depletion experiments, anti-CSF1R antibody (#BE0213, clone AFS98, BioCXell) was administered (400 μg/mouse) thrice a week for two weeks (26,27).

### Immunohistochemistry

Tissues were fixed in 10% paraformaldehyde and embedded in paraffin for sectioning. Sections were deparaffinized by incubation in xylene and rehydrated by sequential incubation in 100%, 95%, 80%, 60% ethanol. The sections were incubated with primary antibody overnight at 4C (Supp Table 1) and subsequently incubated with biotinylated anti-mouse (1:300, Vector Lab) or biotinylated anti-rabbit (1:300, Vector Lab) for 30 minutes at room temperature. Sections were incubated for 30 minutes at room temperature in an ABC reagent (1:300, Vectastain ABC kit, Vector Lab). Sections were developed using 3,3’- Diaminobenzidine (DAB) for 30- 60 seconds, counterstained with hematoxylin, and mounted with Permount. The stained sections were scanned on Digital Pathology Slide Scanner (Philips) and quantified on ImageJ software (NIH).

**Supplementary Table 1:** Antibodies used in Immunohistochemistry.

| Reagent type<br>(species) or resource | Designation | Source or<br>reference | Identifiers | Additional<br>Information |
| --- | --- | --- | --- | --- |
| Antibody | MYC<br>(Rabbit,<br>Monoclonal) | Epitomics | RRID:AB_11000<br>313 | IHC (1:150), IF<br>(1:150), WB<br>(1:1000) |
| Antibody | Phospho<br>Histone3<br>(Rabbit,<br>Polyclonal) | Cell<br>Signaling<br>Technology | RRID:AB_33153<br>5 | IHC (1:100) |
| Antibody | CD4 (Mouse,<br>Monoclonal) | Abcam | RRID:AB_26869<br>17 | IHC (1:100) |
| Antibody | F4/80 (Rat,<br>Monoclonal) | Thermo Fisher | MF48000 | IHC (1:50) |
| Antibody | CD8 (Rat,<br>Monoclonal) | Thermo Fisher | MAI-145 | IHC (1:50) |
| Antibody | PDL1 (Rabbit,<br>Monoclonal) | Cell<br>Signaling<br>Technology | RRID:AB_64988 | IHC (1:100) |

### Mass-cytometry analysis

Infiltration of immune cells in the tumor was assessed using mass-cytometry (CyTOF). After euthanization, the mice were perfused by injecting PBS into the left ventricle. The dissected tumor was cut into small pieces and processed into a single cell suspension using the Tumor Dissociation Kit (MACS Miltenyi) according to the manufacturer’s instructions. The cell suspension was filtered through a 70 µm cell strainer and treated with the ACK lysis buffer (Thermo Fisher). Approximately 5 *10^6^ cells were stained for CyTOF analysis. In brief, the cells were incubated for 15 min at RT with 1 µM Intercalator-Rh (Fluidigm) diluted in Maxpar Cell Staining Buffer (Fluidigm) for dead cell discrimination. Surface Fc receptors were blocked prior to staining by adding 2.5µg Rat anti-mouse CD16/32 Fc block (BD Pharmingen). Without washing out the blocking antibody, the cells were stained with metal-conjugated antibodies according to the list in Supp Table 2 in a total volume of 100 µl per sample and incubated for 45 min at RT. All metal-conjugated antibodies were titrated prior to use. Antibodies were washed out using the Maxpar Cell Staining Buffer and the cells were resuspended in 125 nM Intercalator-Ir (Fluidigm) diluted in Maxpar Fix and Perm Buffer (Fluidigm) and incubated overnight at 4°C. The intercalator was washed out using the Maxpar Cell Staining Buffer following a washing step in Maxpar water (Fluidigm). The cells were kept as a pellet on ice until analysis. Immediately before analysis, the cells were resuspended to a concentration of approx. 0.5 *10^6^ cells/ml in Maxpar water containing 10% (v/v) EQ Four Element Calibration Beads (Fluidigm). The samples were analyzed on the DVS CyTOF 2 instrument (Fluidigm). Data analysis was performed using FlowJo v10 and Cytosplore (42). The cells were first gated on “Live CD45+ single cells” in FlowJo and then tSNE analysis was performed in Cytosplore. The different immune cell populations were defined based on the expression of different markers as shown in Supp Fig 5A. Some antibodies were labeled with metal-conjugates in-house using the Maxpar X8 Multimetal Labelling Kit (Fluidigm) according to the manufacturer’s instructions. For each labeling, 100 µg pure antibody was used, and the metal-conjugated antibody was subsequently resuspended in 120 µl of antibody stabilization buffer (Candor Bioscience, Germany). The metal-conjugated antibodies were titrated prior to use for staining of cells.

**Supplementary Table 2:** Antibodies used in Mass Cytometry.

| **Company:** | **Antigen:** | **Clone:** | **Tag (Mass-Metal):** | **Dilution:** |
| --- | --- | --- | --- | --- |
| Fluidigm | CD45 | 30F11 | 89-Y | 1:200 |
| Fluidigm | Ly-6G | 1A8 | 141-Pr | 1:200 |
| Fluidigm | CD11c | N418 | 142-Nd | 1:100 |
| Fluidigm | CD41 | MWreg30 | 143-Nd | 1:100 |
| Fluidigm | CD115 | AFS98 | 144-Nd | 1:50 |
| Fluidigm | CD4 | RM4-5 | 145-Nd | 1:200 |
| Fluidigm | F4/80 | BM8 | 146-Nd | 1:100 |
| BioLegend | CD44 | IM7 | 147-Sm* | 1:200 |
| Fluidigm | CD11b | M1/70 | 148-Nd | 1:100 |
| Fluidigm | CD19 | 6D5 | 149-Sm | 1:200 |
| Fluidigm | CD24 | M1/69 | 150-Nd | 1:200 |
| Fluidigm | CD64 | X54-5/7.1 | 151-Eu | 1:100 |
| Fluidigm | CD3e | 145-2C11 | 152-Sm | 1:200 |
| Fluidigm | CD274/PD-L1 | 10F.9G2 | 153-Eu | 1:200 |
| Fluidigm | TER-119 | TER-119 | 154-Sm | 1:100 |
| BioLegend | CD133 | 315-2C11 | 155-Gd* | 1:50 |
| R&D Systems | CCR2 | 475301R | 156-Gd* | 1:50 |
| BioLegend | CD103 | 2E7 | 158-Gd* | 1:100 |
| Fluidigm | CD279/PD-1 | 29F1A12 | 159-Tb | 1:100 |
| BioLegend | CD25 | 3C7 | 160-Gd* | 1:50 |
| Fluidigm | CD40 | HM40-3 | 161-Dy | 1:100 |
| Fluidigm | Ly-6C | HK1.4 | 162-Dy | 1:200 |
| BioLegend | CD152/CTLA-4 | UC104B9 | 163-Dy* | 1:200 |
| Fluidigm | CX3CR1 | SA011F11 | 164-Dy | 1:200 |
| BioLegend | CD163 | S15049I | 165-Ho* | 1:50 |
| Fluidigm | CD117/c-kit | 2B8 | 166-Er | 1:100 |
| Fluidigm | CD335/NKp46 | 29A1.4 | 167-Er | 1:200 |
| Fluidigm | CD8a | 53-6.7 | 168-Er | 1:200 |
| Fluidigm | CD206/MMR | C086C2 | 169-Tm | 1:50 |
| Fluidigm | CD169 | 3D6.112 | 170-Er | 1:50 |
| BioLegend | CD69 | H1.2F3 | 171-Yb* | 1:100 |
| BioLegend | CD39 | 24DMS1 | 172-Yb* | 1:50 |
| BioLegend | SiglecF | ES22-10D8 | 173-Yb* | 1:50 |
| BioLegend | IFNAR1 | MAR1-5A3 | 174-Yb* | 1:50 |
| Fluidigm | CD127 | A7R34 | 175-Lu | 1:100 |
| Fluidigm | FceR1a | MAR-1 | 176-Yb | 1:100 |
| Fluidigm | MHC-II | M51/14.15.2 | 209-Bi | 1:400 |

## SUPPLEMENTARY FIGURES

**SUPPLEMENTARY FIGURE 1.**
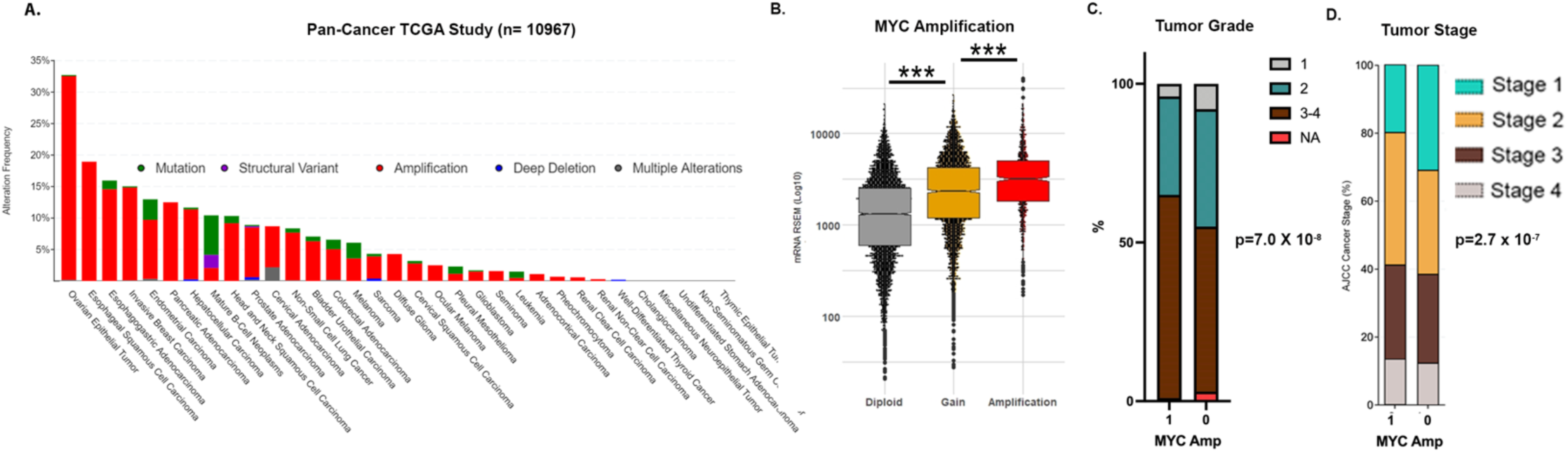
MYC gene amplifications in Human Cancers. A. Prevalence of *MYC* gene amplifications across the 33 TCGA cancers. B. MYC gene amplification strongly correlates with *MYC* mRNA expression across 33 cancers in the TCGA cohort. C. Association of *MYC* amplification in tumors with tumor grade across 33 cancers in the TCGA cohort. D. Association of *MYC* amplification in tumors with tumor stage across 33 cancers in the TCGA cohort.

**SUPPLEMENTARY FIGURE 2.**
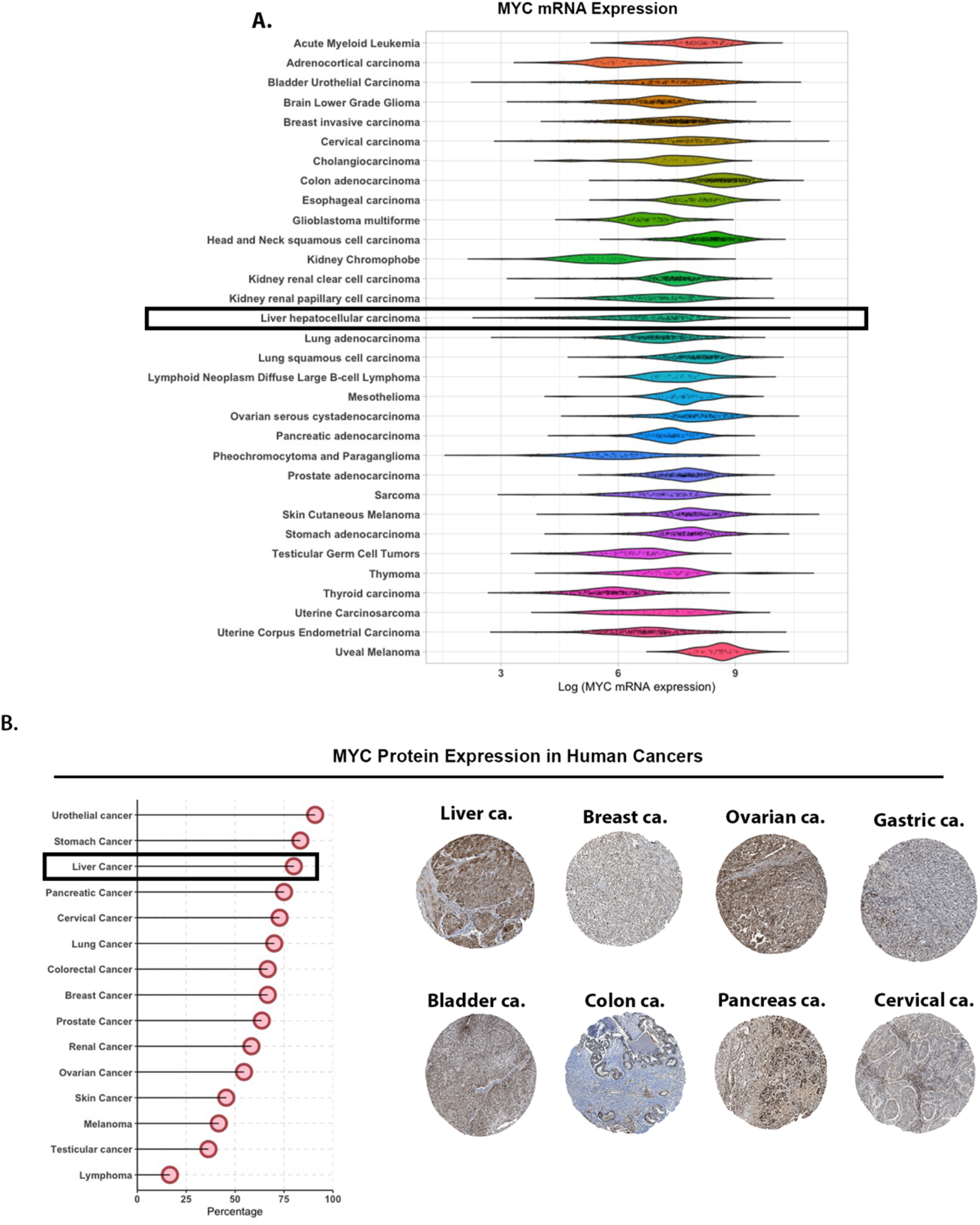
MYC mRNA and Protein Overexpression in Human Cancers. A. *MYC* gene mRNA expression across the 33 TCGA cancers. B. Data from the Human Protein Atlas showing top 10 cancers with MYC protein overexpression (https://www.proteinatlas.org/ENSG00000136997-MYC/pathology).

**SUPPLEMENTARY FIGURE 3:**
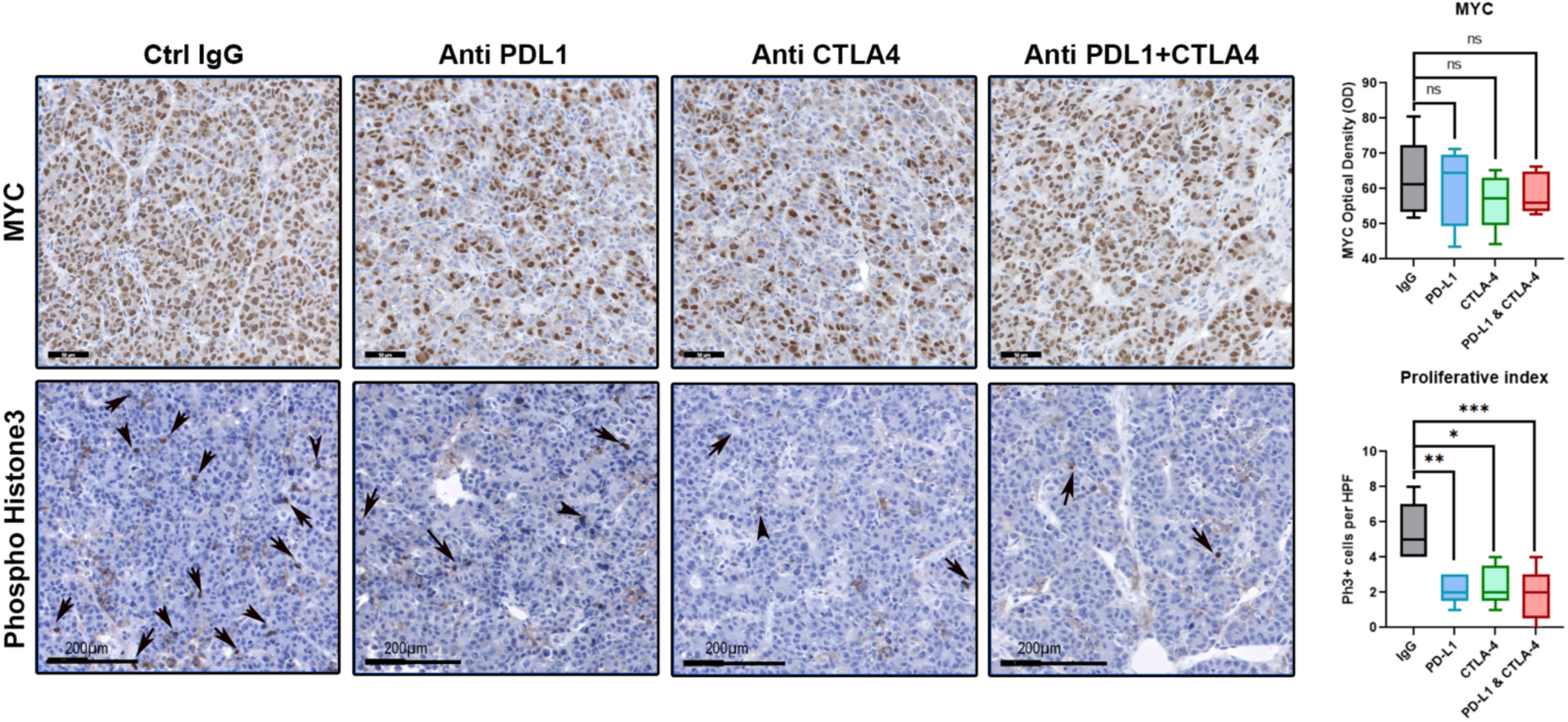
Immune Checkpoint Inhibition Leads to Proliferative Arrest in MYC-HCC. Immunohistochemistry showing MYC and phospho-histone 3 staining and quantification in MYC-HCC treated with IgG control, PDL1 antibody, CTLA4 antibody, or their combination.

**SUPPLEMENTARY FIGURE 4:**
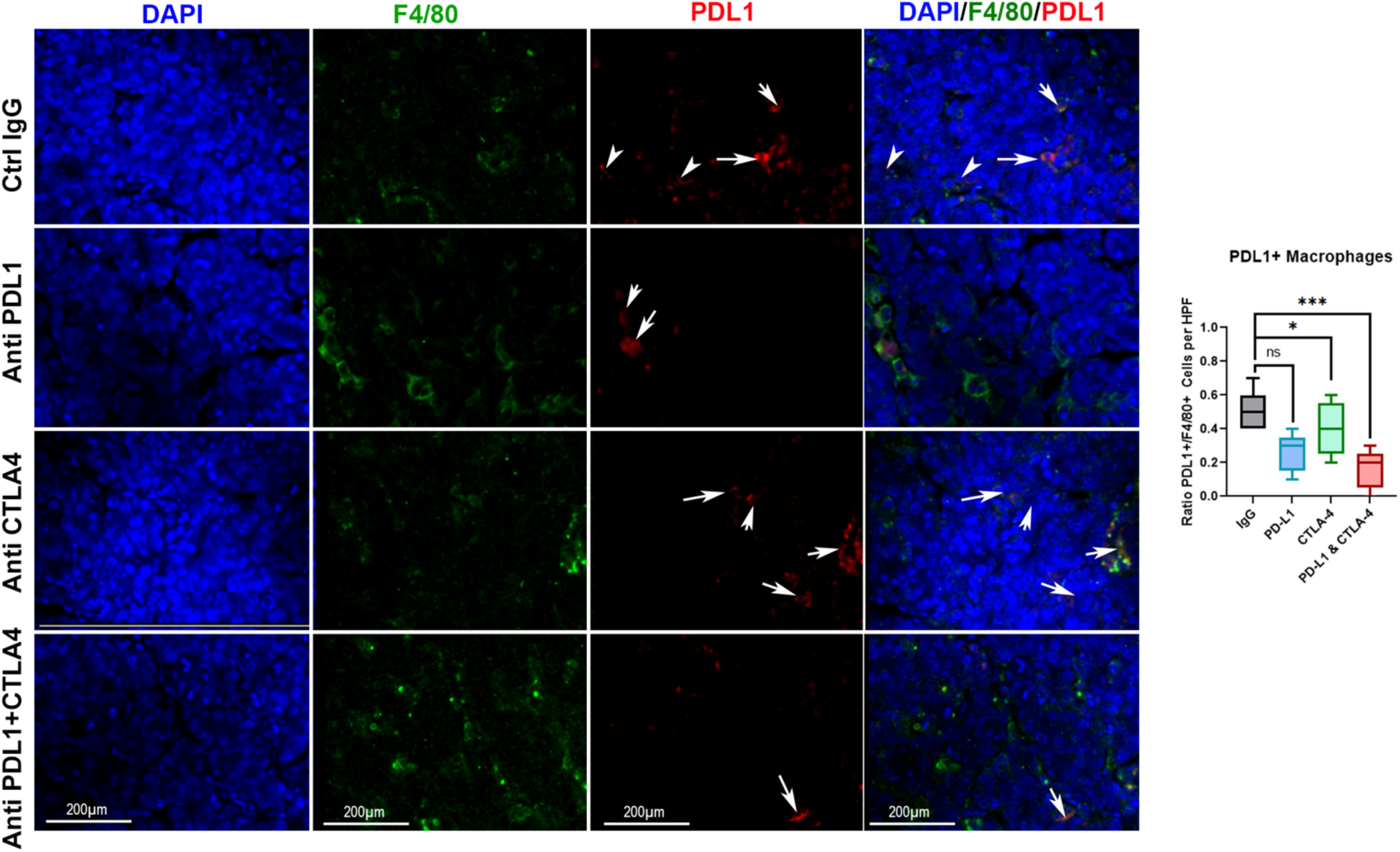
Macrophage repolarization with combination Immune Checkpoint inhibition in MYC-HCC. Immunofluorescence showing PDL1 and F4/80 double staining and quantification in MYC-HCC treated with IgG control, PDL1 antibody, CTLA4 antibody, or their combination.

**SUPPLEMENTARY FIGURE 5:**
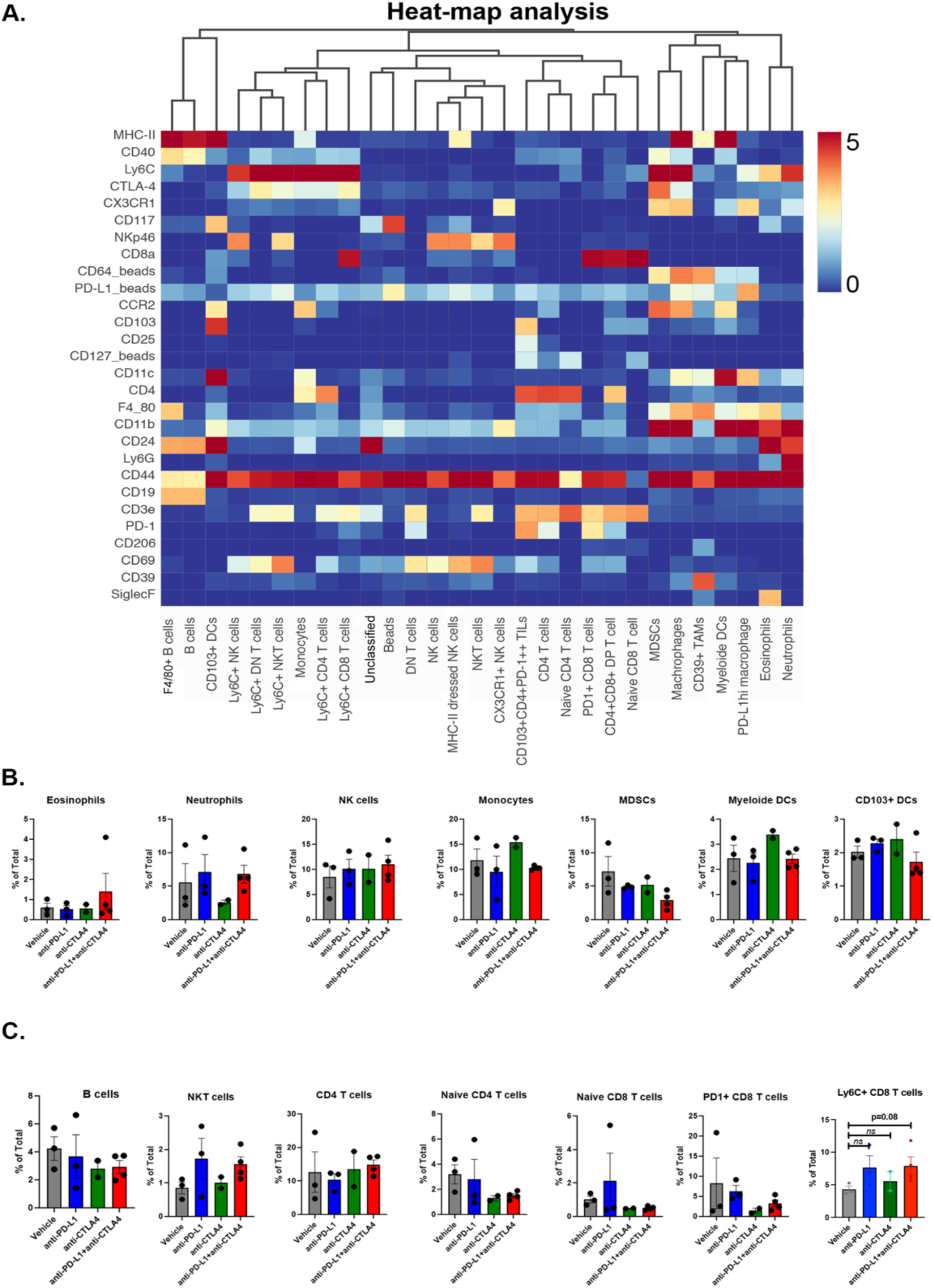
Mass Cytometry Analysis of Immune Response to Immune Checkpoint Inhibitors in MYC-HCC. A. Heatmap shows expression of diffuse immune markers in the identified immune subsets from the mass cytometry analysis of MYC-HCC treated with IgG control, PDL1 antibody, CTLA4 antibody, or their combination. B. tSNE plot shows the myeloid population highlighted in MYC-HCC treated with IgG control, PDL1 antibody, CTLA4 antibody, or their combination. C and D. Comparison of abundance of multiple immune subsets in MYC-HCC treated with IgG control, PDL1 antibody, CTLA4 antibody, or their combination.

**SUPPLEMENTARY FIGURE 6:**
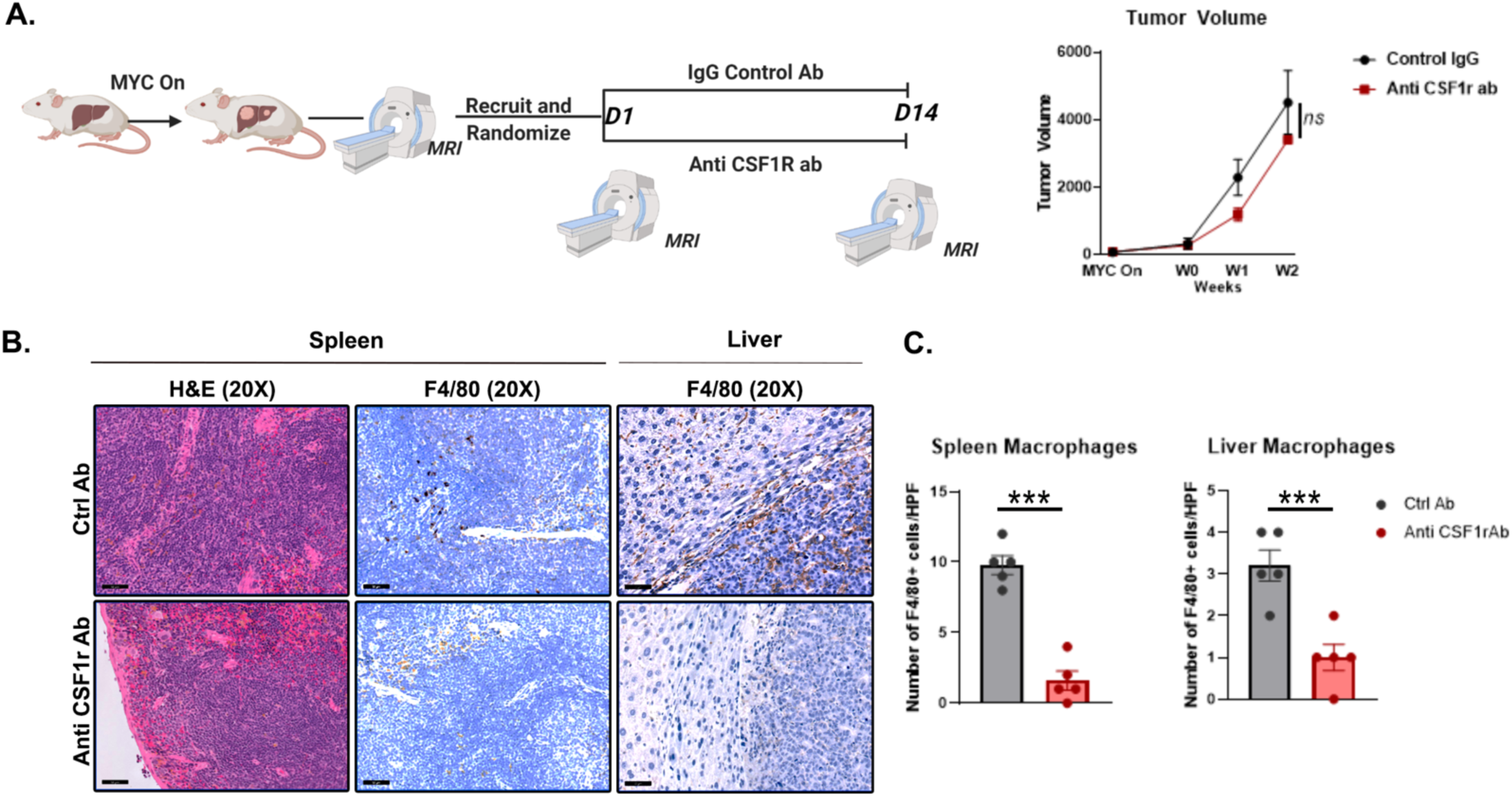
Anti CSF1R Treatment in MYC-HCC. A. Experimental scheme of MYC-HCC mice treated with anti CSF1R antibody or control antibody. No difference in tumor progression was noted on MRI volumetric tumor measurement. B. Representative images of spleen and IHC for F4/80 confirming macrophage depletion with anti CSF1R antibody both in the spleen and the liver.

